# The Spike protein of SARS-coV2 19B (S) clade mirrors critical features of viral adaptation and coevolution

**DOI:** 10.1101/2022.08.12.503822

**Authors:** Bidour K. Hussein, Omnia M. Ibrahium, Marwa F. Alamin, Lamees A.M Ahmed, Safa A.E Abuswar, Mohammed H. Abdelraheem, Muntaser E. Ibrahim

## Abstract

Pathogens including viruses evolve in tandem with diversity in their animal and human hosts. For SARS-coV2, the focus is generally for understanding such coevolution on the virus spike protein since it demonstrates high mutation rates compared to other genome regions, particularly in the receptor-binding domain (RBD).

Viral sequences of the SARS-coV2 19B (S) clade and variants of concern from different continents, were investigated, with a focus on the A.29 lineage which presented with different mutational patterns within the 19B (S) lineages in order to learn more about how SARS-coV2 may have evolved and adapted to widely diverse populations globally.

Results indicated that SARS-coV2 went through evolutionary constrains and intense selective pressure, particularly in Africa. This was manifested in a departure from neutrality with excess nonsynonymous mutations and a negative Tajima D consistent with rapid expansion and directional selection as well as deletion and deletion-frameshifts in the N-terminal domain (NTD region) of the spike protein.

In conclusion, viral transmission during epidemics through population of diverse genomic structure and marked complexity may be a significant factor for the virus to acquire distinct patterns of mutations within these populations in order to ensure its survival and fitness, hence in the emergence of novel variants and strains.

**Importance:** In this study, we justify the fact that the virus’s evolution varies across continents, with each continent showing different amounts and patterns of mutations and deletions, which was manifested in the 19B (S) clade of SARS-coV2, particularly in areas with high population complexity, such as Africa, despite the low rate of sampling and data sharing. The findings show that SARS-coV2 was subject to evolutionary constraints and intense selective pressure. This study will contribute to the scanty amount of research on the SARS-coV2 coevolution and adaptation, in which the host variation is of great significance in understanding the intricacies of viral host coevolution.

## Introduction

SARS-coV-2 is a member of the Coronaviridae family, with wide range of viruses that affect humans, animals, or both (1). Pathogens are being shown to co-evolve in tandem with diversity in animals and humans (2)(3)(4)(5)(6). The exact pathway and timeline of the virus’s emergence and appearance of human cases is still unknown. Many theories exist to track the possible evolution of SARS-coV2 from animals, including recombination events that began with bat corona viruses (RmYN02, RpYN06, and PrC31); it was found to be the closest ancestor for the virus in the whole genome apart from the spike protein in which RaTG13 bat-derived virus is the closest. No intermediate host has been determined so far (2). In the past two years, the virus evolved to give 12 variants, five of them are dominating and each with specific unique set of mutations. These variants include (Alpha (B.1.1.7 and Q lineages), Beta (B.1.351 and descendent lineages), Gamma (P.1 and descendent lineages), Epsilon (B.1.427 and B.1.429), Eta (B.1.525), Iota (B.1.526), Kappa (B.1.617.1), 1.617.3, Mu (B.1.621, B.1.621.1), Zeta (P.2), Delta (B.1.617.2 and AY lineages) and lastly Omicron (B.1.1.529 and BA lineages), which was reported in early November 2021. These variant have been divided into 3 categories either being variants of concern, interest or high consequences, the last which does not include any variant to date (https://www.cdc.gov/coronavirus/2019-ncov/variants/variant-info). Although they circulate the globe, some dominate specific countries. SARS-coV2 mutations are mainly concentrated on the spike protein and Open reading frame1 (ORF1), but as time passed, mutations expanded to include other open reading frames (ORFs) and structural proteins including the Membrane, Envelope and Nucleocapsid proteins.

From the host side, a Spike protein main function is coupling Angiotensin converting enzyme 2 receptor (ACE2), recognizing and fusing to facilitate viral entry to the host cell (7). With the emergence of more transmissible and mutable variants, understanding the evolutionary characteristic of SARS-coV2’s spike genomic region is critical for predicting the path reinfection, vaccination, and therapeutics will take (8).

The spike protein contains several conserved areas but the region of RBD in S1 subunit are the highest mutable region of the spike. S1 is where the initiation of the attachment of the virus to the ACE2 starts (9)(10).

SARS-coV2 N-terminal domain (NTD region) of the spike protein also brought attention because its evolution is related to alteration of the viral antigenicity and promoting immune escaping. The more mutations and/or deletions in this region, the faster SARS-CoV-2 will adapt, evolve and evade the immune system (11).

The mutational robustness found in this virus demonstrates its strength to tolerate host range expansion and adaptation, phenotypic plasticity or environmental stressors such as temperature, virulence or attenuation, antigenicity and immune escape (12). Furthermore, uncontrolled community transmission of SARS-CoV-2 increases the possibility of the emergence of more transmissible variants, which is determined by host diversity in specific countries (13)(14).

Here we investigate SASRS-coV2 sequences specifically the19B (S) clade, the first dominating variant after the ancestral virus of Wuhan (https://clades.nextstrain.org/), obtained from various countries and continents in order to understand the sequel of evolutionary processes and viral adaptation in disparate environments particularly the putative effect of population variation on its distribution and mutational variations.

## Methodology

### Study Design

This is a retrospective cross sectional study to address the global variation in the SARS-coV2 evolution, and how it may have adapted to the host selective pressure. This analysis for the 19B (S) clade, the first dominating variant after the ancestral virus of Wuhan, covering the period from the start of the pandemic until February 2022.

### Viral Genome Sequences

A total of 15,537 viral sequences of 19B (S) clade were procured from GISAID EpiCoV database (https://www.gisaid.org/) for the different continents at the period from the start of the pandemic until February 2022. Sequences of the lineages from A to A30, Bat, Pangolin as well as the variants of concern including Alpha, Beta, Gamma, Delta, Lambda and Omicron were also downloaded from the same database. They were chosen according to their first appearance in the collection dates. More focus was on the 19B (S) lineage A.29 that has been downloaded also.

We filtered sequences for the study based on the length of the sequence not to be less than 29,000 nucleotides, percentage of gaps, N stretches (unidentified nucleotides) of less than 5%, lack of clusters of mutations, and overlapping of reading frames. It was evaluated using GISAID EpiCoV database and NEXTSTRAIN web tool (https://clades.nextstrain.org/).

### Sequences Analysis

Sequences were analyzed using the Next Generation Sequencing (NGS) analysis packages targeting areas of variations in all sequences included in the analysis, COVID-19 genome annotator online tool for annotation (http://giorgilab.unibo.it/coronannotator/) and the NEXTSTRAIN online tool for defining clades for each sequence. Sequences were aligned to SARS-coV 2 isolate Wuhan-Hu-1 with the accession number NC 045512.2S and mutations were examined. The variations of nonsynonymous and synonymous mutations were estimated for the whole genome and the spike protein for each sample.

### Phylogenetic analysis

Phylogenetic analysis was carried out for those variants using the MEGAX software (https://www.megasoftware.net/)(15). Maximum likelihood analysis was performed with bootstrapping (1000 iterations).

### Evolutionary distance and Neutrality Testing

The evolutionary distance and neutrality were tested by Tajima neutrality test using MEGAX software. *P* value were estimated for the proportion of synonymous to nonsynonymous as indicator of neutrality according to Kimura (16).

### 19B (S) A.29 lineage analysis

The lineage of 19B (S) clade A.29, was taken as an example showing different pattern of mutations. Using a Python script, the frequency of mutations were calculated for each country involving the whole genome. Countries with reported A.29 were from Africa (Gambia and Sudan), Asia (India and Jordan), Europe (United Kingdom, Germany, and Belgium) as well as Oceania (Australia) and North America (the United States of America and Canada).

### Data visualization

Frequency of mutations for the lineage A.29 was displayed on bar charts and tables. It was displayed also in a time line chart. Secondary RNA structures for the spike protein were then constructed by first extracting the Spike protein region using the seqkit command (https://bioinformaticsworkbook.org/) and then uploading the sequences to the RNAFOLD online tool (http://rna.tbi.univie.ac.at/cgi-bin/RNAWebSuite/RNAfold.cgi). For secondary structures created, ensemble diversity and Minimal Free Energy (MFE) were estimated. To show the variation in the structure of the spike protein and the NTD region, a 3D model was created using the EXPASY translation (https://web.expasy.org/translate/) and the SWISSMODEL online tools (https://swissmodel.expasy.org/interactive).

### Comparison with human sequences

A simple diagram was plotted depicting the possible relationship between effective population size and variation in some genetic markers in the host with the virus based on published data on GISAID EpiCoV database and published human variation data (17)(18)(19)(20).

## Results

The distribution of the 19B (S) clade samples varied, expectedly, between and within countries due to the wide differences in the sequencing efforts. Findings revealed the existence of different patterns of distribution of the 19B (S) clade lineages across continents, in which for example, the A.23 and A.27 were dominating Africa with approximate percentages of 39.67% and 14.996% respectively, while the A Lineage dominated Asia with 61.1%. The A.2 found to be common in Europe, North America, South America and Oceania with approximate percentages of 37.23%, 40.623%, 76.23% and 56.93% respectively. In Africa, the lineages A18 to A30 found to be more common than in other continents (Supplementary table 1).

In Africa, the total number of the 19B(S) clade samples is 1147, the nonsynonymous to synonymous mutations were 1468\1013, deletions were 28 and deletion-frameshifts were 18 across the whole genome. Whereas in North America where n = 8313, it has 3822 nonsynonymous mutations, 2225 synonymous, 55 for both deletions and deletions-frameshifts (Table 1).

**Table 1:**
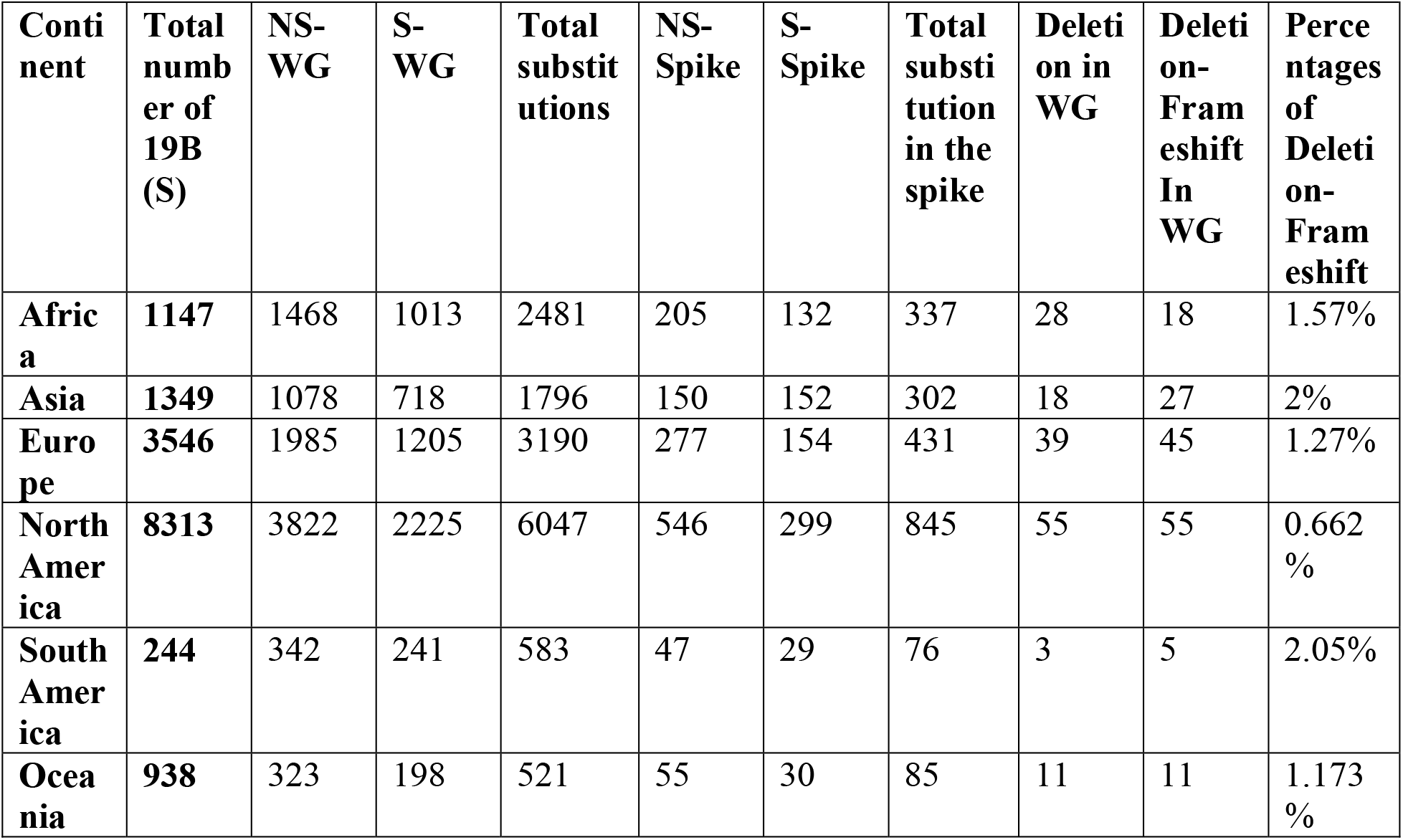
The distribution of the 19B (S) clade in the different continents and the variation in the nonsynonymous, synonymous, deletion and deletion frameshift. Both the whole genome and the spike alone were presented. Abbreviations: NS-WG: nonsynonymous mutations in the whole genome, S-WG: synonymous mutations in the whole genome, NS-Spike: nonsynonymous mutations in the spike protein and S-Spike: synonymous mutations in the spike protein. Note: Not all countries has updated sample uploading in the GISAID platform.

The same pattern of variation applies to other lineages and variants of concern. Based on the dates of collection reported in the databases combined with phylogenetic analysis, 11 of the lineages were found to have originated most likely in Africa including Senegal (A.11), Serra Leone (A.12), Burkina Faso (A.18), Côte d’Ivoire (A.19), Mali (A.21), Uganda (A.23), Kenya (A.25), Niger (A.27), Egypt (A.28), Gambia (A.29), and finally Angola (A.30). Nine of the other sub variants originated in Asia and the rest were scattered in Europe and North America (Figure 1).

**Figure 1:**
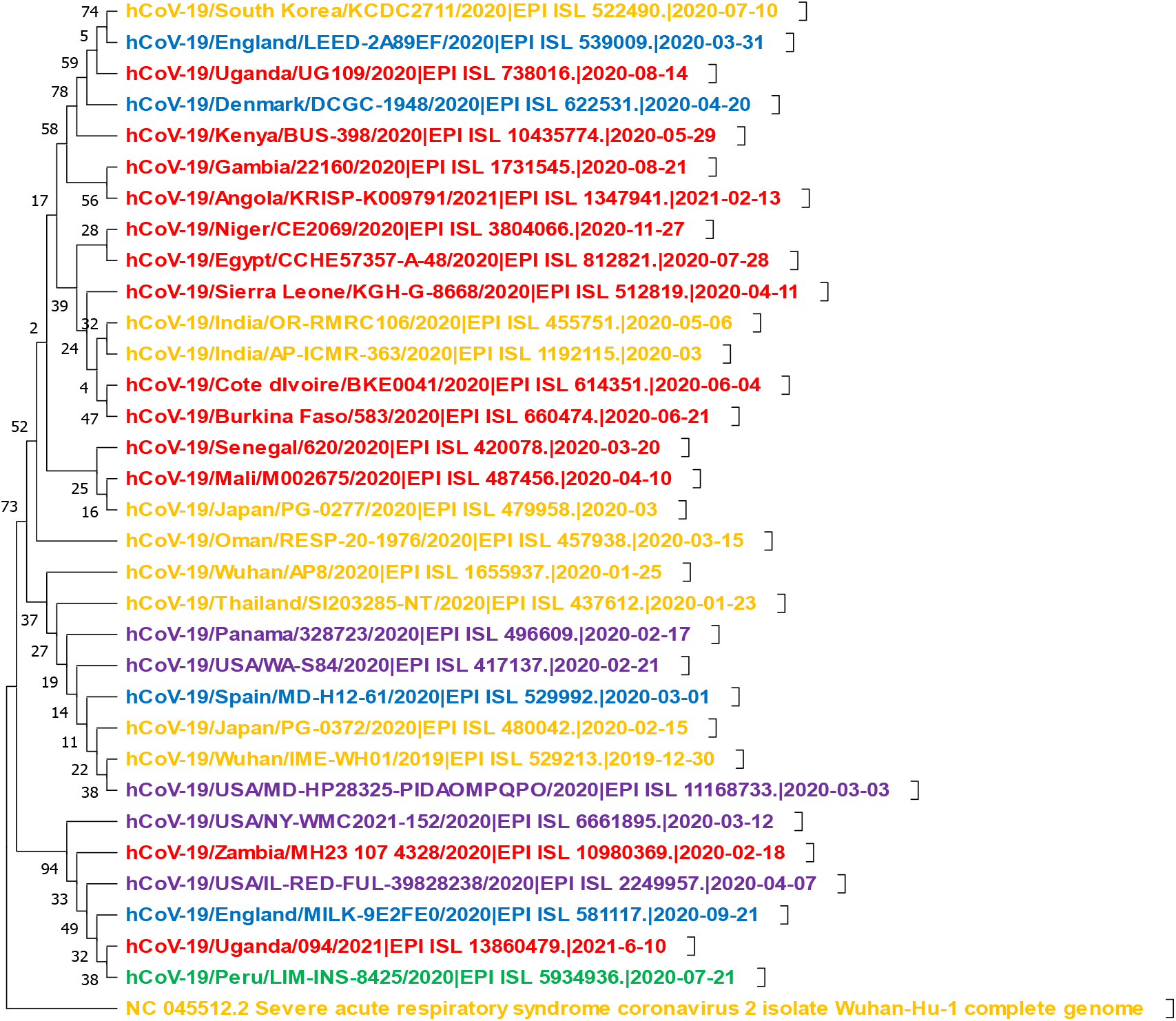
Evolutionary analysis by Maximum Likelihood method. The evolutionary history was inferred by using the Maximum Likelihood method and Kimura 2-parameter model. The tree with the highest log likelihood (-45061.28) is shown. The percentage of trees in which the associated taxa clustered together in 1000 bootstraps is shown next to the branches. Initial tree(s) for the heuristic search were obtained automatically by applying Neighbor-Join and BioNJ algorithms to a matrix of pairwise distances estimated using the Maximum Composite Likelihood (MCL) approach, and then selecting the topology with superior log likelihood value. This analysis involved 33 nucleotide sequences. Note: colores indicate, Red (Africa), Yellow (Asia), Blue (Europe), Purple (North America) and Green (South America).

Signals of selection were manifested in the excess of nonsynonymous mutations in the global sample x235.7. (P=0.0001), particularly in the African continent in comparison to other continents (Z=3.91 P=0.0001) despite the small sample size. Tajima’s Neutrality test returned a negative D value of -2.646764 consistent with rapid expansion and directional selection.

The A.29 lineage of the 19B (S) clade presented with specific mutational pattern and hence was selected to address questions of adaptation and coevolution. These samples had shared mutations that were different from other sub variants in the 19B (S) clade in which it include: 15 nonsynonymous mutations, 9 synonymous mutations, 2 deletions, one frameshift and extra-genic mutation. Frequencies of those mutations are shown in (Figure 2), (Supplementary figure 1, and 2). Deletions and deletion-frameshift were concentrated in the NTD region of the S1 subunit of the spike protein whereas nonsynonymous mutations were scattered in different ORFs of the viral genome including the spike protein itself. In the 3’ UTR of the genome, there were extra-genic mutations at Coronavirus 3’ stem-loop II-like motif (s2m; located in the region from 29,728-29,768) at position 29,742 and 29,739 (Supplementary figure 3).

**Figure 2:**
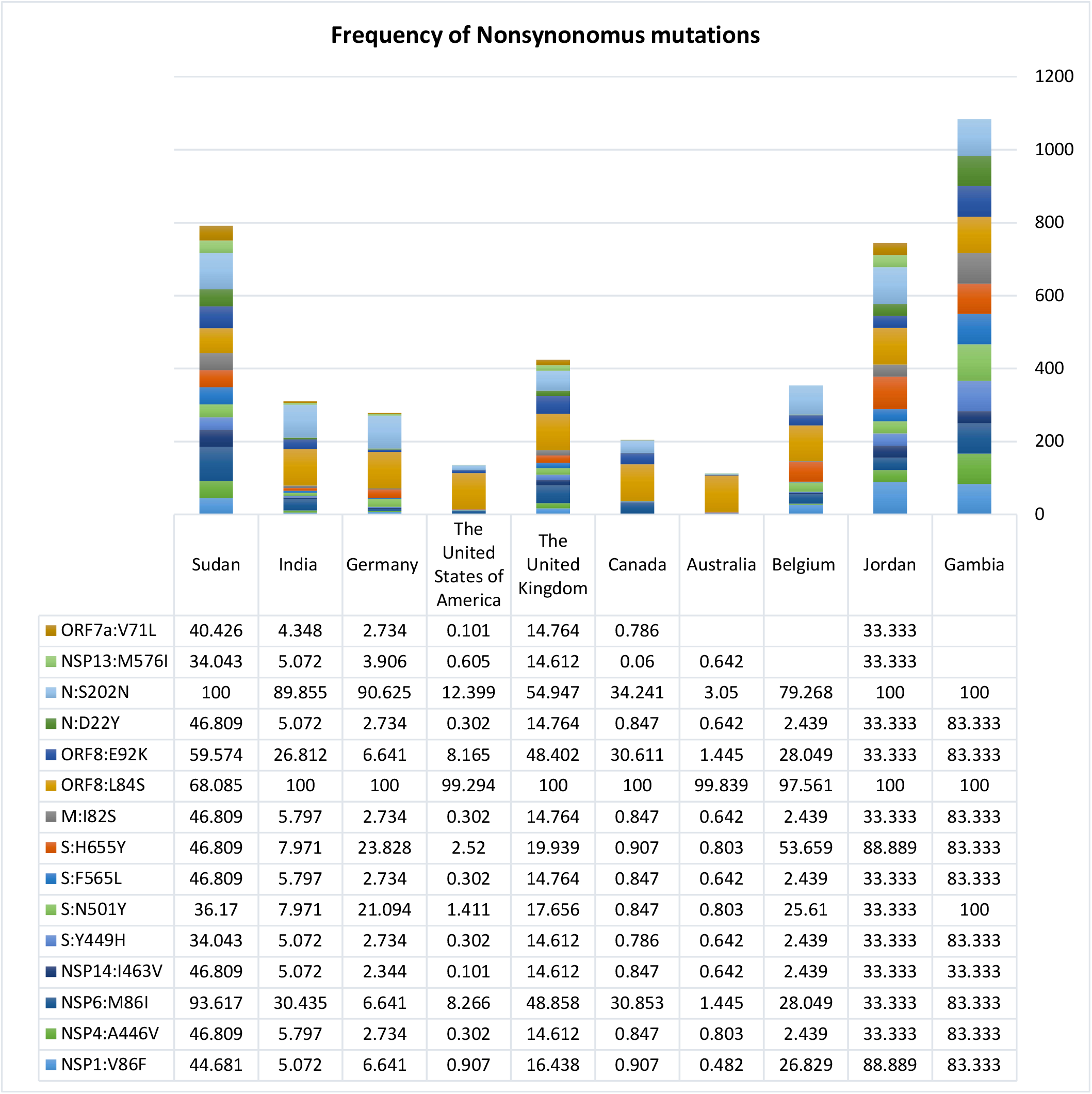
The frequency of the shared nonsynonymous mutations in samples of 19B (S) clade A.29 lineage with different pattern among countries over the whole genome include: N:S202N, N:D22Y,ORF8:E92K, ORF8:L84S, M:I82S, S:H655Y, S:F565L, S:N501Y, S:Y449H, NSP14:I463V, NSP6:M86I, NSP4:A446V, NSP1:V86F.

Based on the CoVariants website (https://covariants.org/) which gives an overview of SARS-CoV-2 variants and their mutations which is supported by data from GISAID platform mutations from above mentioned samples were found mainly in those variants of concern as a nonsynonomus mutations including: S:N501Y dominating 20I (Alpha, V 1), 20H (Beta, V2) and 20J (Gamma,V3), S:H655Y dominating 20J (Gamma,V3), 21K(Omicron) and 21L(Omicron), M:I82S dominating 21B (Kappa), S:V143 and S:N211 dominating 21K(Omicron), and finally the Extra-genic mutation in the 3’ UTR:29,742 in 21A, I and J (Delta) and 21B (Kappa) as G nucleotide substituted to T which is synonymous mutation, but in those samples it was substituted with an A nucleotide. None of the known variants showed substitution at 3’ UTR: 29,739, which changed from C to T. Visualization of structural variations in the spike protein were presented in secondary and 3D structures, all in comparison to the reference genome of SARS-coV2, Bat-derived virus of Yunnan, and Pangolin. There were small variation from the reference genome in the minimal free energy and ensemble diversity in secondary structures and multiple areas of deletions were demonstrated in the 3D structures (Supplementary figure 4, 5, 6, 7, 8, 9, and 10).

Those samples were scattered across the pandemic years. The majority appeared in 2021, with the exception of samples from Kombo city in Gambia and Kassala city in Sudan, samples reported in August and December 2020 respectively (Figure 3).

**Figure 3:**
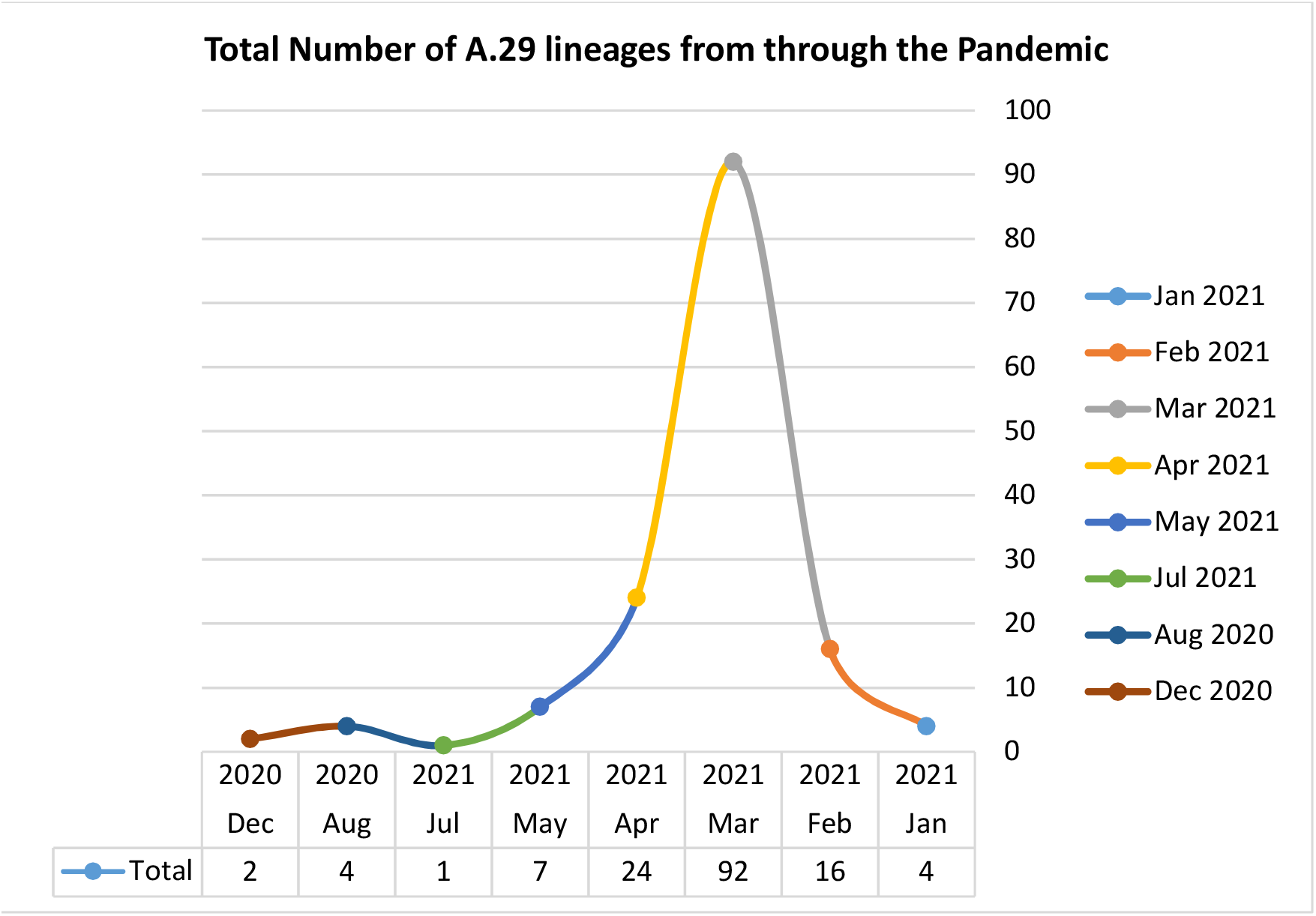
Illustrates the spread of samples of the A.29 lineage across countries over the two-year pandemic. The first sample was reported in Kombo Gambia on August 2020 n=four, followed by Sudan in December 2020 in Kassala n=two.

As the human-COVID paired samples are scanty or non-existent to allow addressing the potential relationship between the virus and the host diversity, human genetic data was obtained from the literature and depicted in a simple diagram. The figure shows a crude correlation between the high genetic variation of the host like in Africa and the virus population diversity in the same continent. The genetic marker R1b1b2 and Omicron are an example of such correlations (Figure 4) (https://www.gisaid.org/)(17)(18)(19)(20).

**Figure 4:**
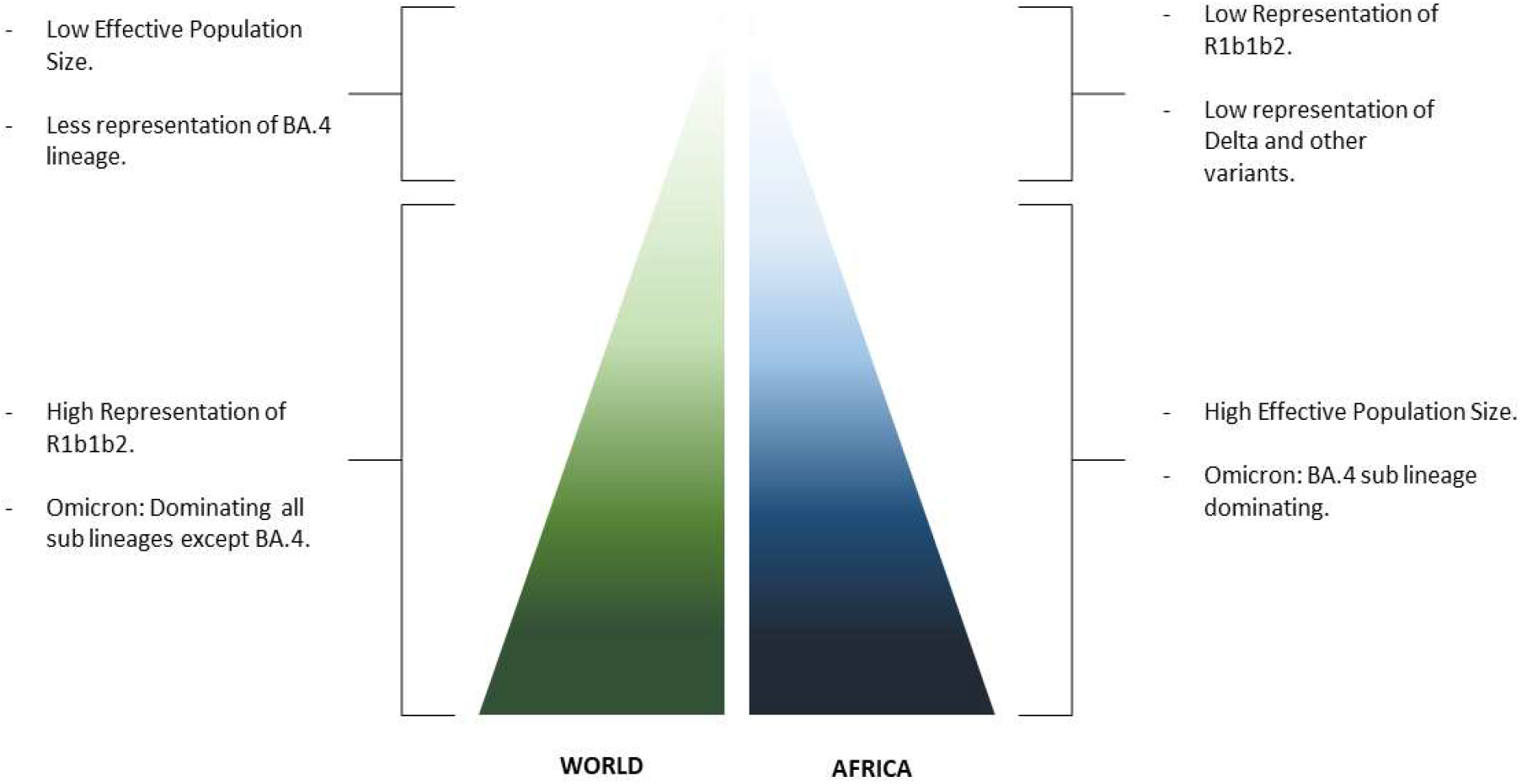
A schematic representation of how variation in the host genome and population size are correlated with viral variation, as seen in the sub-lineage of Omicron variant distribution based on multiple studies and the amount of SNP and deletions in both virus and host. Here, The high effective population size and the presence of the suceptibility marker R1b1b2 in low frequency in Africa could be one of the related causes for viral variant dominancy.

## Discussion

For a virus like SARS-coV2, to jump the species barriers, it needs to pass through stages of natural selection and incuring more genomic changes to inrease its affinity to human ACE2. This was applied to the strucure of the spike protein of the Bat-derived virus of Yunnan and Pangolin samples which showed multiple deletions and deletion framshifts not represented in the referance genome of SARS-coV2 (21). Viruses are able to evolve within the host itslef resulting in varients and strains that could infect and transmit better (22). Once this happens in certain population, a variant will form clusters and allow for more transmission and escaping of the immune surveillances of the host. Still in countries like in Africa where a highly diverse populations exist (17)(18)(19), the dynamics and time line for the virus evolution and adaptation is not well understood due to the lack of adequate sampling. Taking Sudan as an example in which the 19B (S) clade was dominating and represented with different pattern of mutations in high frequency (49%) could be a clue for the virus track to adapt with population variability in order to increase its survival (21)(23). Example of some mutations found is the H655Y substitution which enhances spike cleavage and viral growth (24). N501Y and Y449H substitutions, which are believed to increase the virus transmisabilty and fittness, although it has been shown that the Y449H substitution alone lead to decrease in the affinity of binding to the ACE2 recepter, but when it came along with the N501Y it was spedculated that they will enhance binding affinity and escaping, based on epistatic shifting and modulation (25). In addition there are the deletion and deletion frameshift in the NTD region, which plays an important role in conformation of the spike protein structure, binding to the ACE2 and immune escaping, although it shows higher rate of conservation (26). These mutational patterns are scattered in different variants and strains of concern and interest, rather than being grouped in one variant including the 19B (S) strain. A pattern of similar mutations were shown in a few other countries including gambia, India, Jordan, the United Kingdom, Germany, Belgium, the United States of America, Canada, and Australia with low frequencies, possibly due to migration, since the first samples were detected in Gambia in August followed by Sudan in December in the year 2020 based on collection dates reported in GISAID platform.

Deletions could be either deleterious, and the virus start reverting to its ancestral state or compensatory with changes that could mask the effect of the pervious mutations. Such mechanism will guarantee recovery of fitness, all depending on the effective population size of the organism. The mechanism is common in the RNA viruses (27). For the virus to have deletion-frameshift followed by deletion, means that the virus is going through an intense selective pressure and through a survival strategy to overcome variation, which has been shown in Omicron and Delta variants that manifest such breadth of deletions in the S1 subunit (28). By comparing SARS-CoV2 to the SARS-CoV-2-related coronavirus (SC2r-CoV) lineages from bats and pangolins, researchers found out that NTD area gained indels during viral transmission across animals are the same as those reported during human transmissions (7). According to one theory, increasing global human population immunity to natural SARS-CoV-2 infection is driving such a large selection pressure on the virus genome, which may select for convergent deletions at NTD to avoid being neutralized by neutralizing antibodies against NTD (29). Another study demonstrated that, rather than working in antibody avoidance mechanism, NTD deletion in certain locations boosts viral infectivity by boosting the incorporation of cleaved S into virion (13).

As RNA virus, SARS-coV2 could affect the host as a quasispecies population hence for indels mutation it can be maintained which will not be the case if the infected host came as one clonal population with the same genome sequences (30)(31)(32). In addition, these quasispecies are maintained after selection and bottleneck events (33).

Having an adaptive landscape will allow overcoming the high functional constrains of the virus for more transmissibility and infectivity (27). However, this landscape should not be from the pathogen side alone. A hypothesis called the Red Queen mentioned that host and pathogen reciprocally struggle to maintain constant levels of fitness (34), and virus copy number and mutational rate is highly determined by the variability within the host (35)(36)(28). Many studies where done in coadaptation of different ancient viruses, taking Adenovirus lineages as an example, it appeared to have speciated and coevolved with different vertebrate families and host shifts to new taxa were accompanied by changes in genome content and those non-coevolved showed more severe pathology (35). Others studied how this relationship can affect different part of the human genome, for example the mitochondrial DNA, hepatitis C virus is able to affect reversibly the diversity of mitochondrial DNA, in which it can be used as a biomarker to investigate the stage of infection(36). Other studies mentioned that having some genetic markers represented in certain population has a large effect on the disease progression, for example, the representation of Y chromosome ancestry marker R1b1b2 in certain countries including in Africa with low frequencies, is associated with a lower mortality rate from COVID 19 infection (20).

Interestingly, African viral sequences, appeared to be ancestral to most of the lineages of the 19B (S) clade and for two variants of concern including Beta and Omicron, making the continent together with Asia, the site for the emergence of these variants and in tally with the hypothesis that places of high effective population is probably the main source of novel variants. Tajima’s Neutrality test D value is consistent with rapid expansion and directional selection a major feature of the pandemic. In numerous studies it was discovered that Africans have higher nucleotide and haplotype diversity than non-Africans (5)(17)(18)(19). In another study, ACE2 receptor are presented with some different nonsynonymous mutations in African poulations which were identified as signatures of selection affecting variation at regulatory regions related to ACE2 expression (37).

The dearth of viral sequnces from the African continent in this manuscript might be related to paucity of sampling due to a poorly resourced and ailing health systems or perhaps to the relatively low disease burden. Circumventing such hurdles is essential for a lucid understanding of the epidemiology of the pandemic and in verfying concepts like the above (38)(39).

Finally and in conclusion, we believe that for the virus to evolve within complex populations characterized by a higher effective population size, it may be pivotal for the virus to acquire distinct patterns of mutations that ensures its survival and fitness, thus becoming a source of novel variants of concern something that has been noticed early on in the 19 (S) clade of SARS-coV2. These variations raises many questions related to the function of the mutations and structural motif within the host that may lead to better viral fitness or transmissibility.

## Acknowledgments

The completion of this work would not have been possible without the participation and assistance of a large number of people. Their contributions are gratefully acknowledged and appreciated. However, we would like to express our gratitude to: Dr. Shahd Osman, Mudathir Salih, Muiz M. Hamza, Ahmed Berir, and Mahmoud Koko for their kind assistance.

## Data availability

Sequences are available at the Global Initiative for Sharing All Influenza Data (GISAID EpiCoV database) portal.

## Supplementary tables and Figures

**Supplementary table 1:**
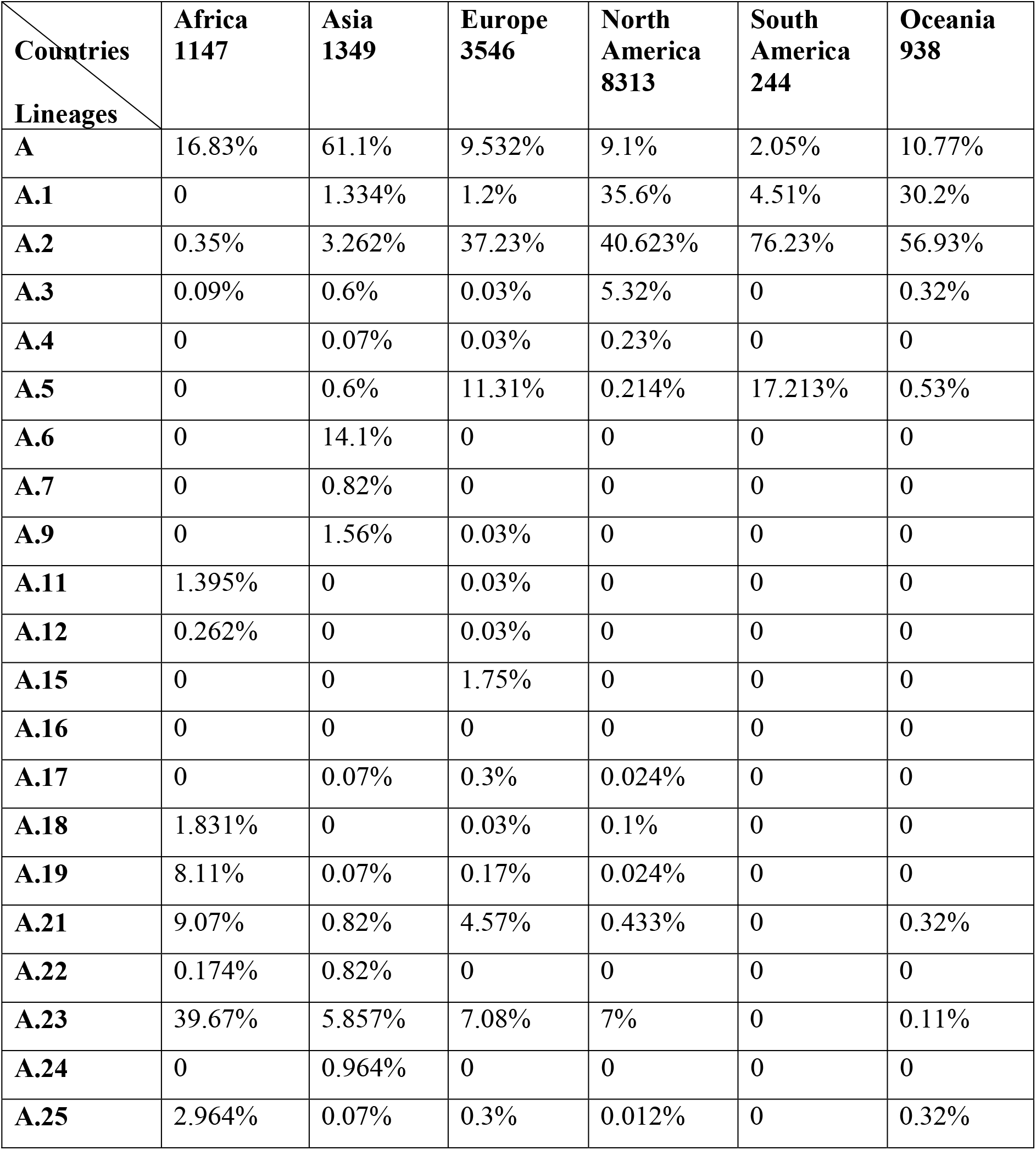
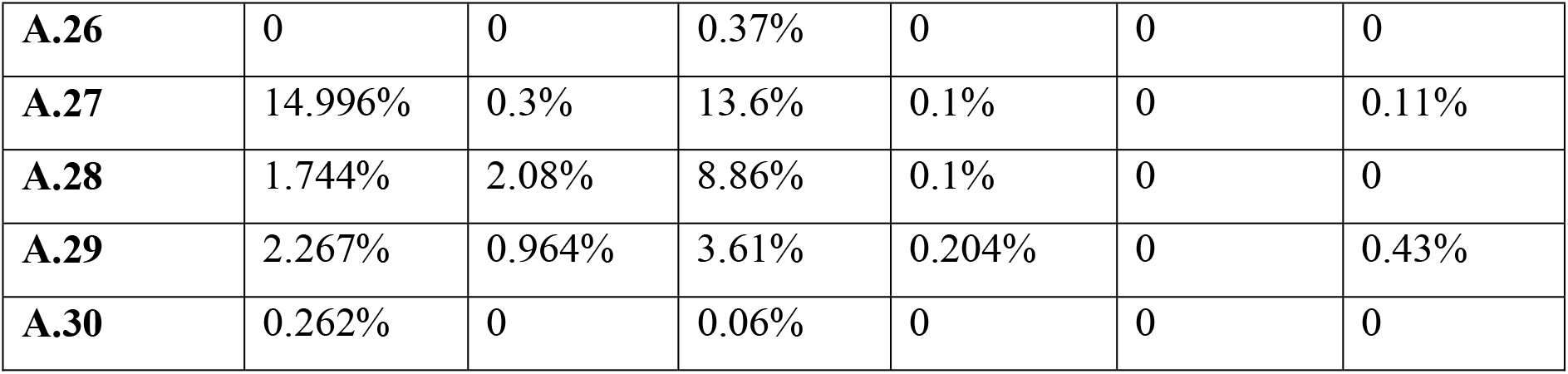
The distribution in the 19B (S) lineages in the different contents, in which the A.23 and A.27 were dominating Africa with approximate percentages of 39.67% and 14.996% respectively, while the A Lineage dominated Asia with 61.1%. The A.2 found to be common in Europe, North America, South America and Oceania with approximate percentages of 37.23%, 40.623%, 76.23% and 56.93% respectively.

| **Countries**<br>**Lineages** | **Africa**<br>**1147** | **Asia**<br>**1349** | **Europe**<br>**3546** | **North**<br>**America**<br>**8313** | **South**<br>**America**<br>**244** | **Oceania**<br>**938** |
| --- | --- | --- | --- | --- | --- | --- |
| **A** | 16.83% | 61.1% | 9.532% | 9.1% | 2.05% | 10.77% |
| **A.1** | 0 | 1.334% | 1.2% | 35.6% | 4.51% | 30.2% |
| **A.2** | 0.35% | 3.262% | 37.23% | 40.623% | 76.23% | 56.93% |
| **A.3** | 0.09% | 0.6% | 0.03% | 5.32% | 0 | 0.32% |
| **A.4** | 0 | 0.07% | 0.03% | 0.23% | 0 | 0 |
| **A.5** | 0 | 0.6% | 11.31% | 0.214% | 17.213% | 0.53% |
| **A.6** | 0 | 14.1% | 0 | 0 | 0 | 0 |
| **A.7** | 0 | 0.82% | 0 | 0 | 0 | 0 |
| **A.9** | 0 | 1.56% | 0.03% | 0 | 0 | 0 |
| **A.11** | 1.395% | 0 | 0.03% | 0 | 0 | 0 |
| **A.12** | 0.262% | 0 | 0.03% | 0 | 0 | 0 |
| **A.15** | 0 | 0 | 1.75% | 0 | 0 | 0 |
| **A.16** | 0 | 0 | 0 | 0 | 0 | 0 |
| **A.17** | 0 | 0.07% | 0.3% | 0.024% | 0 | 0 |
| **A.18** | 1.831% | 0 | 0.03% | 0.1% | 0 | 0 |
| **A.19** | 8.11% | 0.07% | 0.17% | 0.024% | 0 | 0 |
| **A.21** | 9.07% | 0.82% | 4.57% | 0.433% | 0 | 0.32% |
| **A.22** | 0.174% | 0.82% | 0 | 0 | 0 | 0 |
| **A.23** | 39.67% | 5.857% | 7.08% | 7% | 0 | 0.11% |
| **A.24** | 0 | 0.964% | 0 | 0 | 0 | 0 |
| **A.25** | 2.964% | 0.07% | 0.3% | 0.012% | 0 | 0.32% |
| **A.26** | 0 | 0 | 0.37% | 0 | 0 | 0 |
| **A.27** | 14.996% | 0.3% | 13.6% | 0.1% | 0 | 0.11% |
| **A.28** | 1.744% | 2.08% | 8.86% | 0.1% | 0 | 0 |
| **A.29** | 2.267% | 0.964% | 3.61% | 0.204% | 0 | 0.43% |
| **A.30** | 0.262% | 0 | 0.06% | 0 | 0 | 0 |

**Supplementary figure 1:**
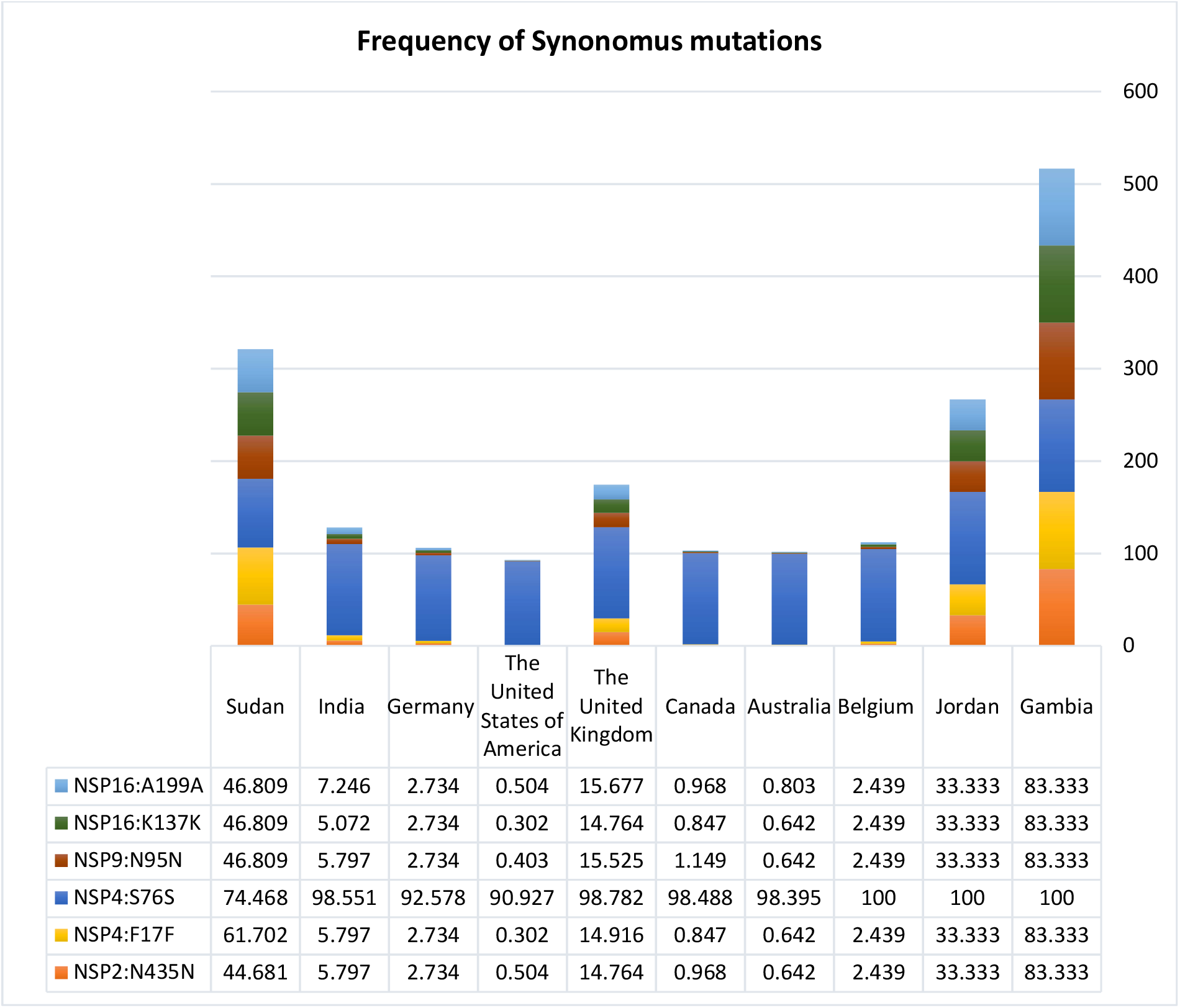
The frequency of the shared synonymous mutations in samples of 19B (S) clade A.29 lineage with different pattern among countries over the whole genome include NSP16:A199A, NSP16:K137K, NSP9:N95N, NSP4:S76S, NSP4:F17F, and NSP2:N435N.

**Supplementary figure 2:**
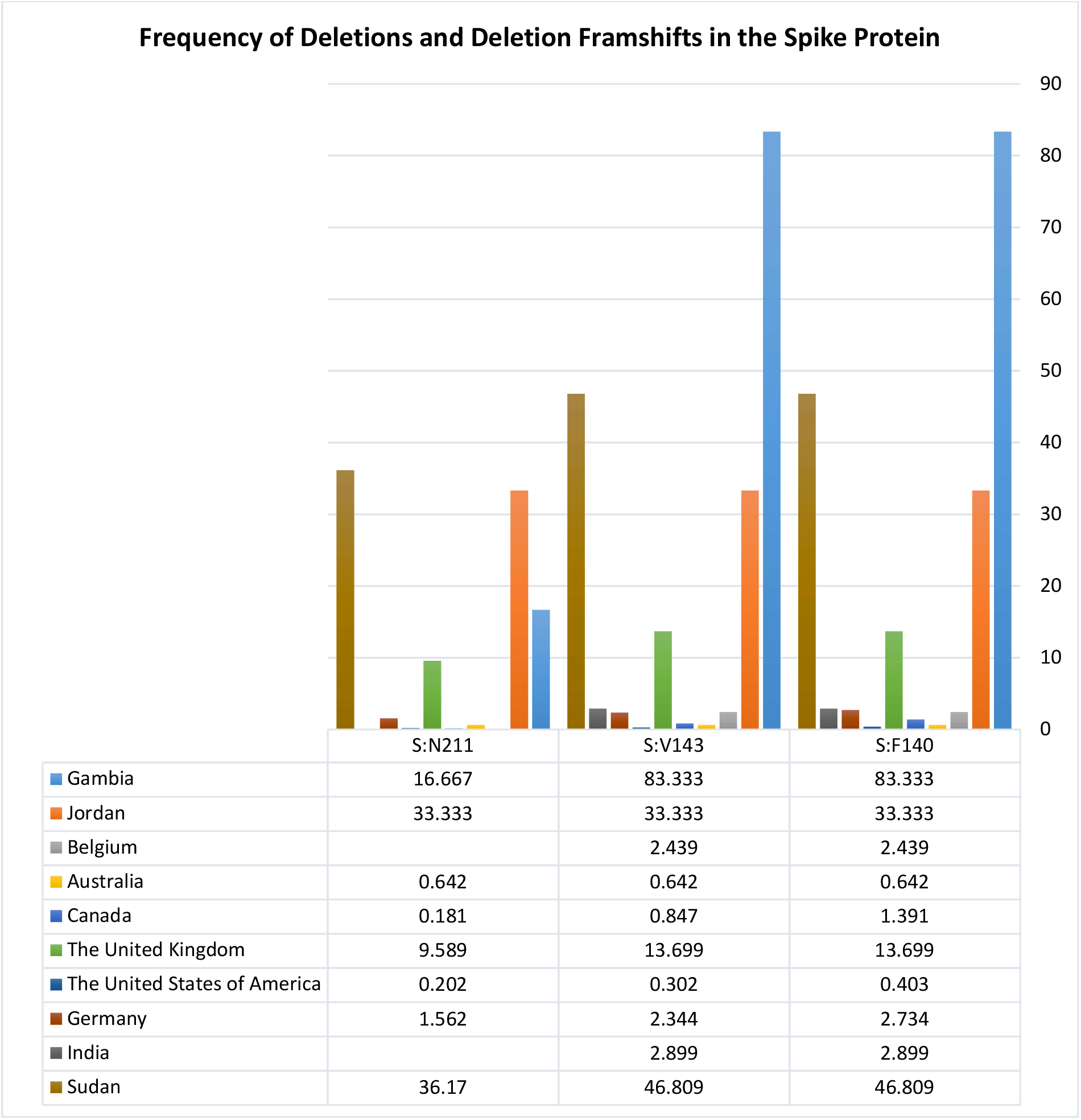
The frequency of the shared Deletion and Deletion-Frameshifts in samples of 19B (S) clade A.29 lineage with different pattern among countries over the whole genome include S: F140, S: V143, S: N211.

**Supplementary figure 3:**
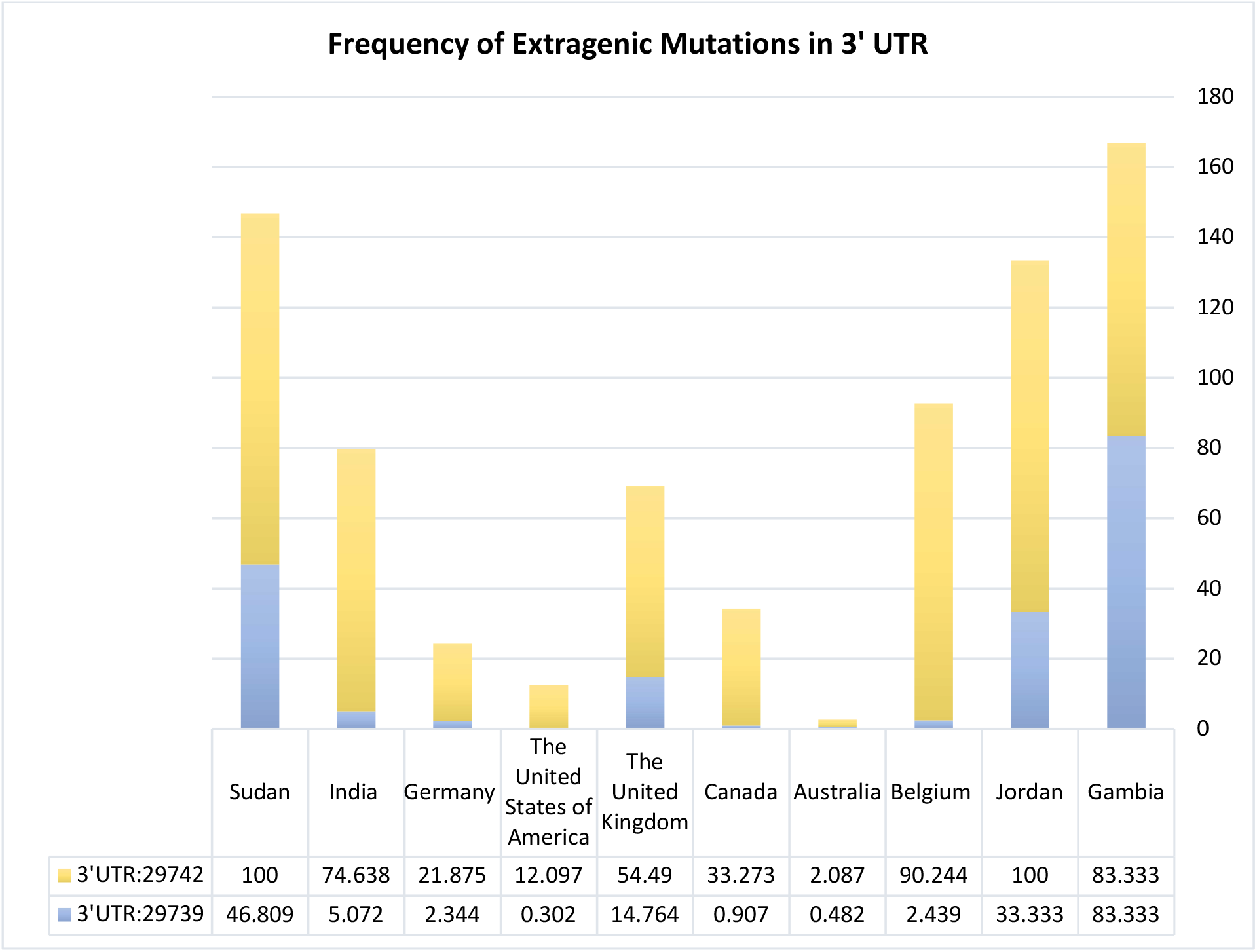
The frequency of shared Extragenic mutations in the 3’UTR region in samples of 19B (S) clade A.29 lineage with different pattern among countries over the whole genome.

**Supplementary figure 4:**
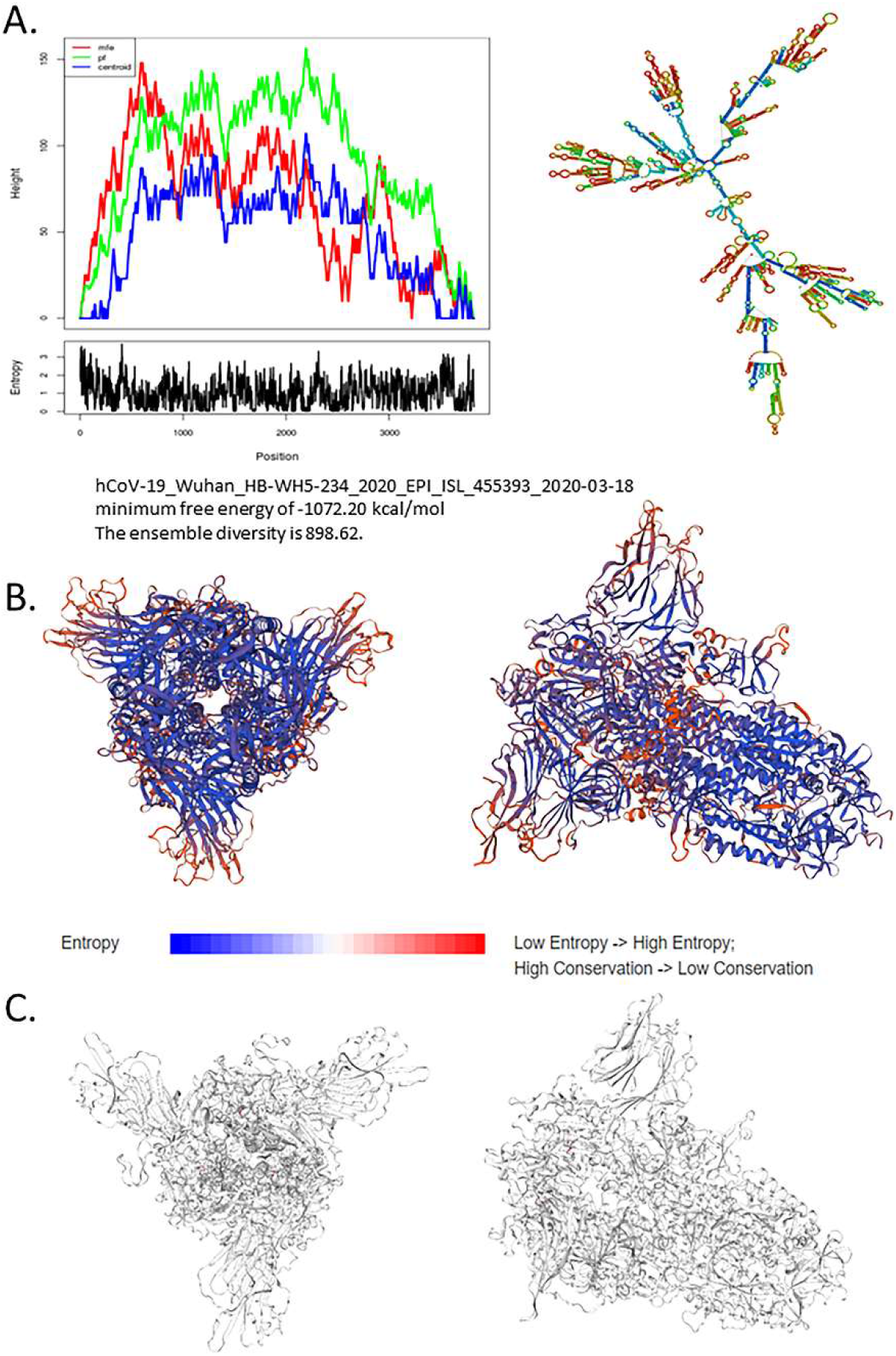
**A**. Secondary structure, for the sample hCoV-19_Wuhan_HB-WH5-234_2020_EPI_ISL_455393_2020-03-18 (Reference: accession number NC 045512.2S), Minimum free energy of -1072.20 kcal/mol, the ensemble diversity is 898.62. **B**. 3D structure of the spike protein using the template 7cn8.1.A, seq identity: 91.68% with the description of Glycoprotein Cryo-EM structure of PCoV_GX spike glycoprotein. **C**. show no deletion in the spike protein.

**Supplementary figure 5:**
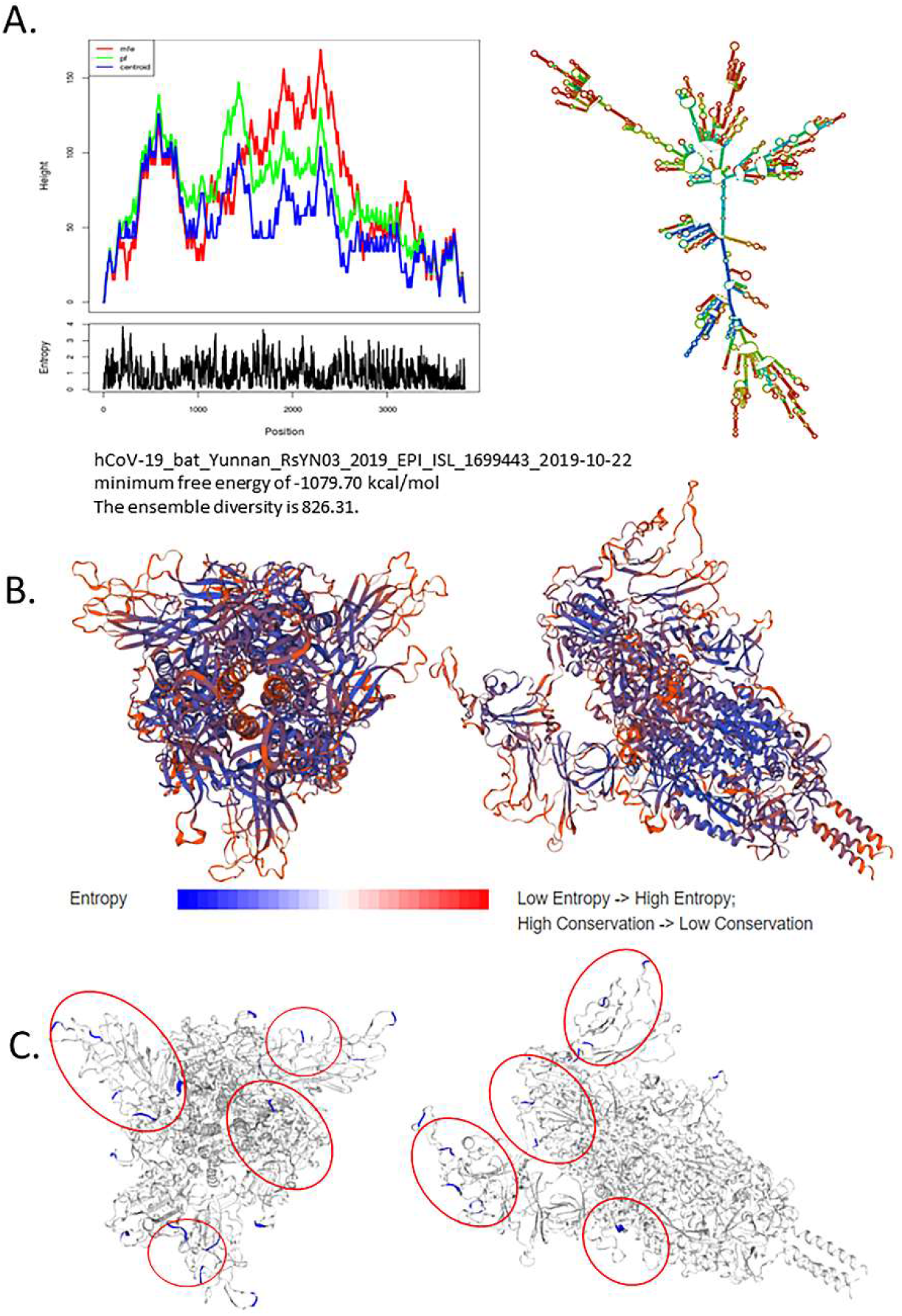
**A**.Secondary structure for the sample hCoV-19_bat_Yunnan_RsYN03_2019_EPI_ISL_1699443_2019-10-22, Minimum free energy of - 1079.70 kcal/mol, and the ensemble diversity is 826.31. **B**. 3D structure of the spike protein using the template 7sbo.1.A, seq identity: 78.48% with the description of Glycoprotein 1 RBD-up 2 of pre-fusion SARS-CoV-2 Delta variant spike protein. **C**. shows multiple deletions in the spike protein marked in blue.

**Supplementary figure 6:**
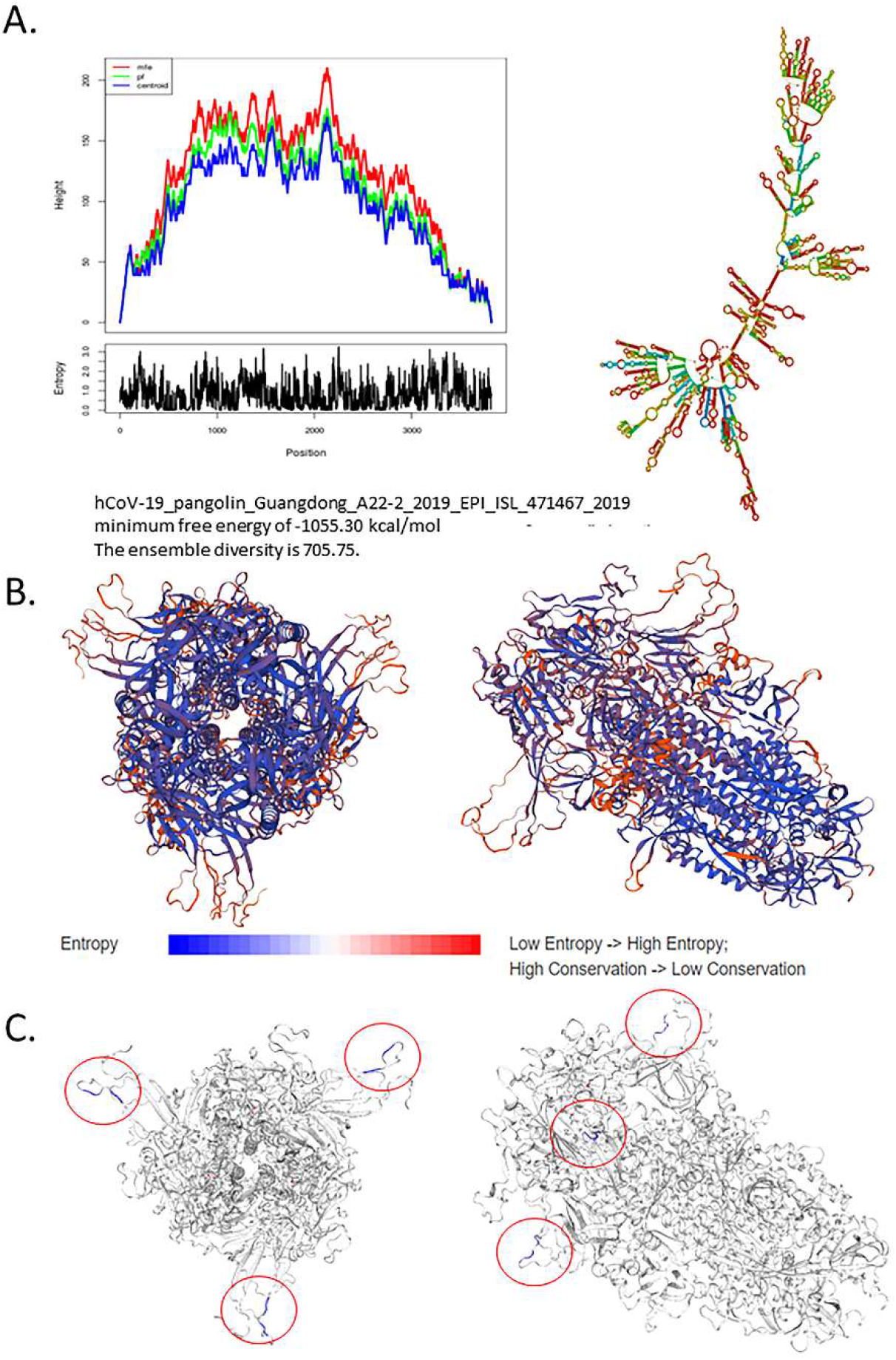
**A**.Secondary structure for the sample hCoV-19_pangolin_Guangdong_A22-2_2019_EPI_ISL_471467_2019, Minimum free energy of - 1055.30 kcal/mol, and the ensemble diversity is 705.75. **B**. 3D structure of the spike protein using the template 7cn8.1.A, seq identity: 90.73% with the description of Glycoprotein Cryo-EM structure of PCoV_GX spike glycoprotein. **C**. shows multiple deletions in the spike protein marked in blue.

**Supplementary figure 7:**
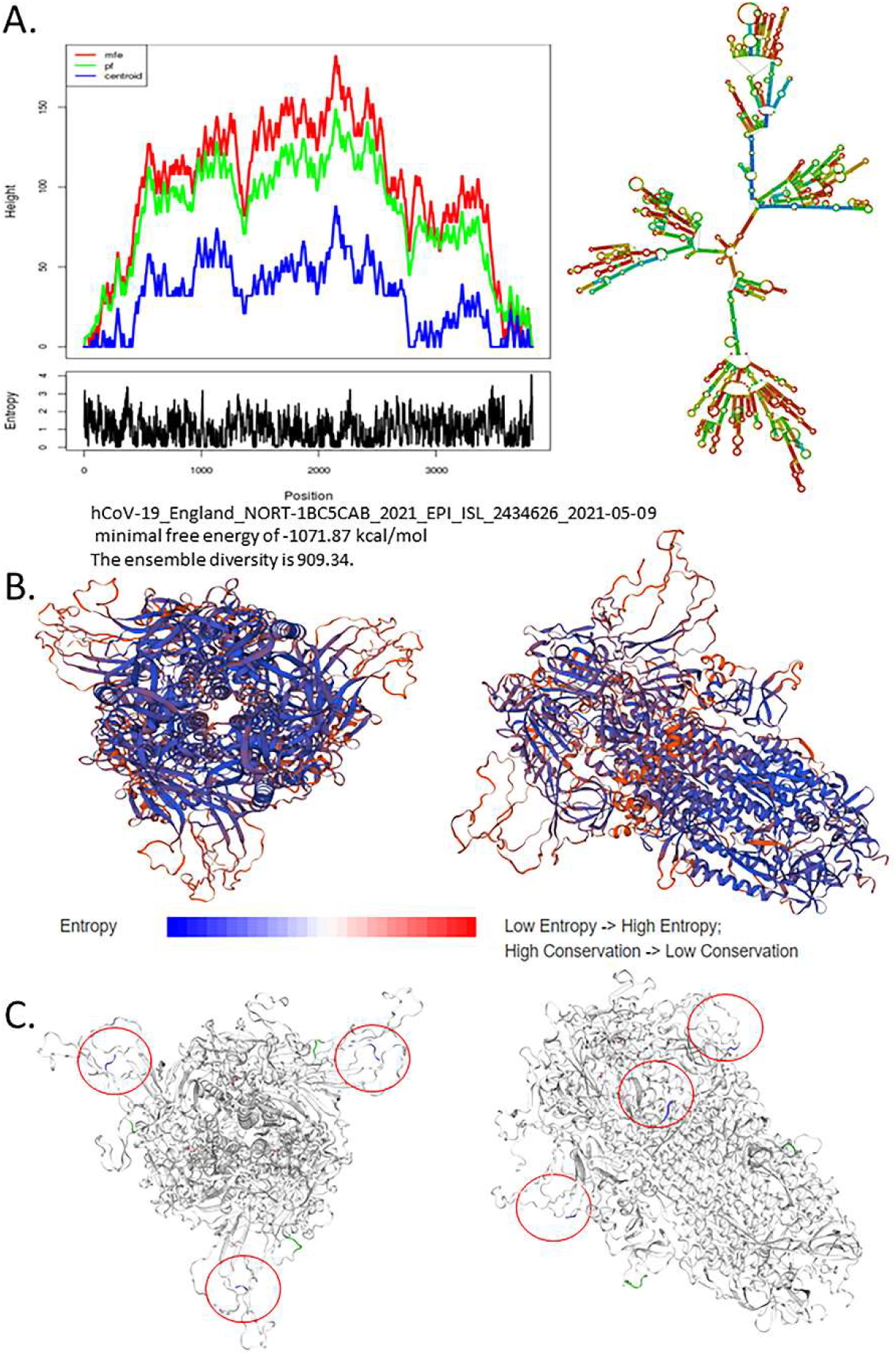
**A**.Secondary structure for the sample hCoV-19_England_NORT-1BC5CAB_2021_EPI_ISL_2434626_2021-05-09, Minimal free energy of -1071.87 kcal/mol, and the ensemble diversity is 909.34. **B**. 3D structure of the spike protein using the template 7cn8.1.A, seq identity: 92.18% with the description of Glycoprotein Cryo-EM structure of PCoV_GX spike glycoprotein. **C**. shows multiple deletions in the spike protein marked in blue.

**Supplementary figure 8:**
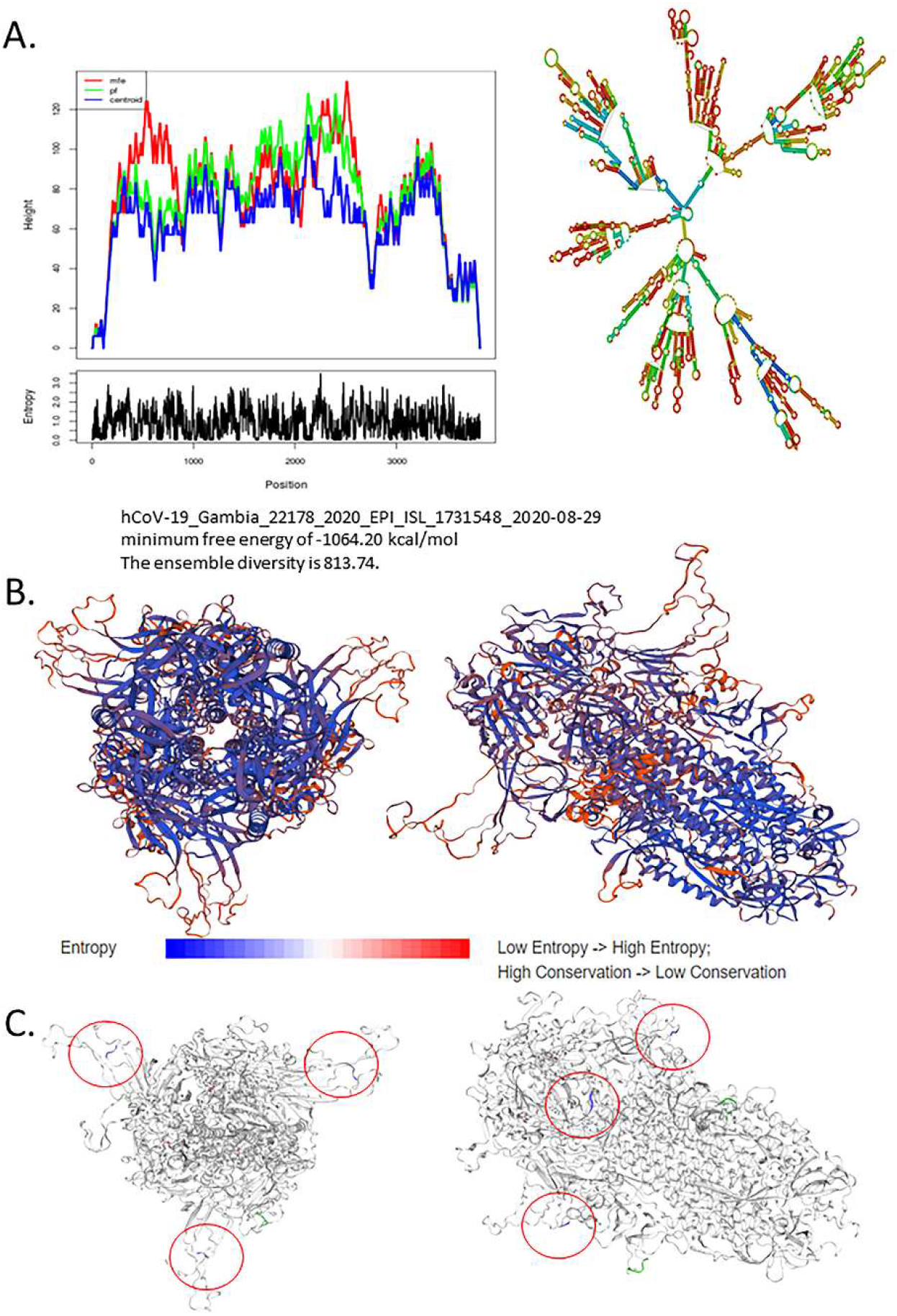
**A**.Secondary structure for the sample hCoV-19_Gambia_22178_2020_EPI_ISL_1731548_2020-08-29, Minimum free energy of - 1064.20 kcal/mol, and the ensemble diversity is 813.74. **B**. 3D structure of the spike protein using the template 7cn8.1.A, seq identity: 92.20% with the description of Glycoprotein Cryo-EM structure of PCoV_GX spike glycoprotein. **C**. shows multiple deletions in the spike protein marked in blue.

**Supplementary figure 9:**
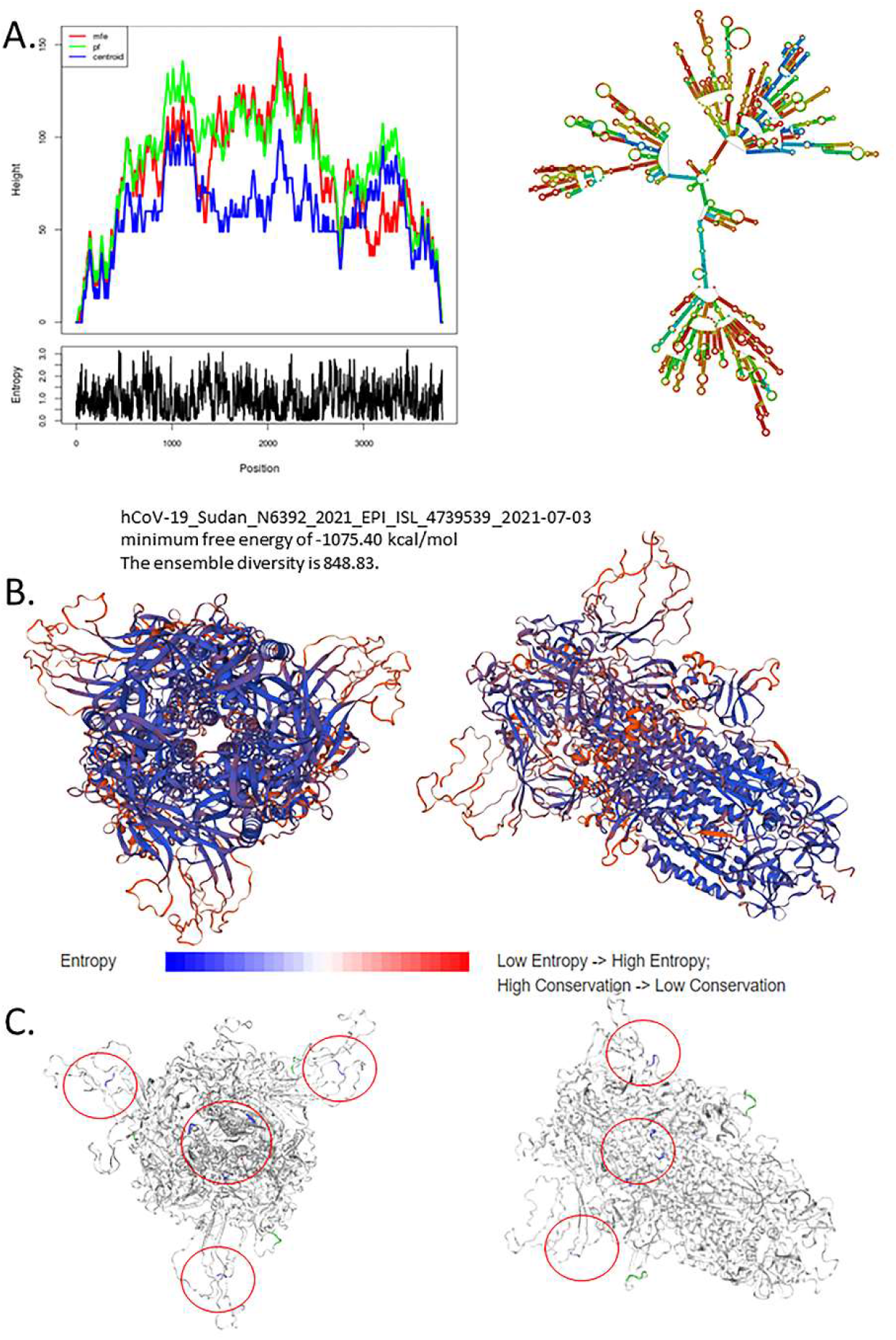
**A**.Secondary structure for the sample hCoV-19_Sudan_N6392_2021_EPI_ISL_4739539_2021-07-03, Minimum free energy of - 1075.40 kcal/mol, and the ensemble diversity is 848.83. **B**. 3D structure of the spike protein using the template 7cn8.1.A, seq identity: 92.08% with the description of Glycoprotein Cryo-EM structure of PCoV_GX spike glycoprotein. **C**. shows multiple deletions in the spike protein marked in blue.

**Supplementary figure 10:**
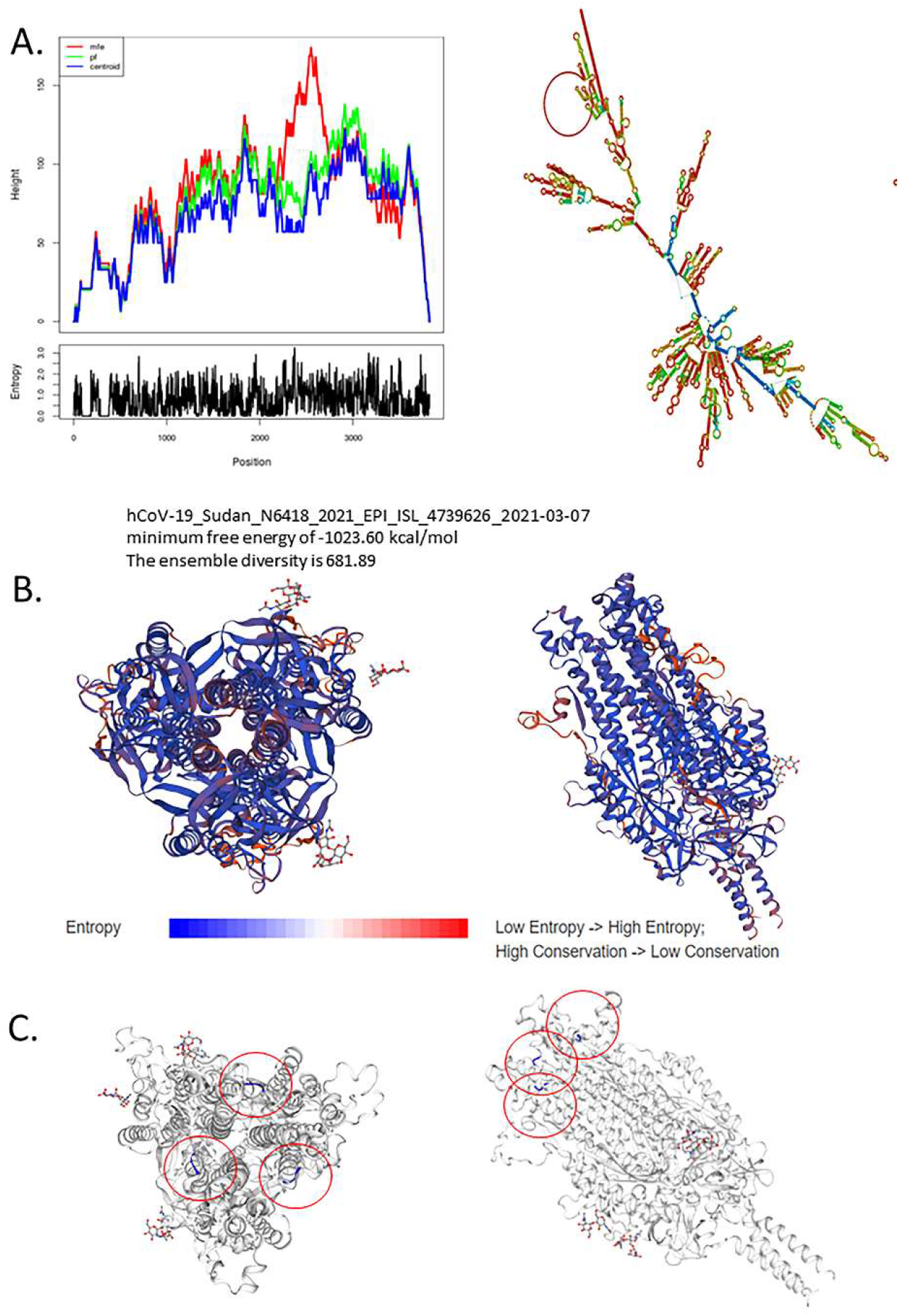
**A**.Secondary structure for the sample hCoV-19_Sudan_N6418_2021_EPI_ISL_4739626_2021-03-07, Minimum free energy of - 1023.60 kcal/mol, and the ensemble diversity is 681.89. **B**. 3D structure of the spike protein using the template 7krs.1.A, seq identity: 100.00% with the description of Spike glycoprotein Structural impact on SARS-CoV-2 spike protein by D614G substitution. **C**. shows multiple deletions in the spike protein marked in blue.

